# AbBERT: Learning Antibody Humanness via Masked Language Modeling

**DOI:** 10.1101/2022.08.02.502236

**Authors:** Denis Vashchenko, Sam Nguyen, Andre Goncalves, Felipe Leno da Silva, Brenden Petersen, Thomas Desautels, Daniel Faissol

## Abstract

Understanding the degree of humanness of antibody sequences is critical to the therapeutic antibody development process to reduce the risk of failure modes like immunogenicity or poor manufacturability. We introduce AbBERT, a transformer-based language model trained on up to 20 million unpaired heavy/light chain sequences from the Observed Antibody Space database. We first validate AbBERT using a novel “multi-mask” scoring procedure to demonstrate high accuracy in predicting complementary determining regions—including the challenging hypervariable H3 region. We then demonstrate several uses of AbBERT at various points along the antibody design process. AbBERT enhances in silico antibody optimization via deep reinforcement learning by utilizing its learned embeddings as additional observations during optimization. Within a larger computational antibody design platform, AbBERT has been successfully applied as an additional design objective, where it displays strong correlations with computational tools predicting antibody structural stability. Finally, mutant antibody sequences that have been scored as unfavorable by AbBERT have shown corresponding low yields when expressed in cells. These use cases demonstrate the power of language modeling within computational antibody design.

## Introduction

While enormous strides in synthetic biology have allowed a scientist to digitally specify the sequence of amino acids that fully describe a designed protein, transmit this to a commercial provider, and have that object reliably made and delivered, it remains difficult to predict what properties the final protein may have and how those properties result from the original design. A promising alternative to directly predicting properties is to take masses of readily available protein sequence data and learn the implicit structure in these sequences, under the hypothesis that more conformant protein sequences will also be more performant. Both very general approaches, attempting to cover all protein classes, and special-purpose approaches, targeting particular, high-value protein classes, are appropriate. In this work, we apply language modeling techniques to the primary sequences of one of the most important protein classes, antibodies.

Under the hypothesis above, a self-supervised model of antibody sequences could be leveraged to inform downstream design decisions and contribute to a better understanding of properties such as the antibody’s safety as a therapeutic or its ease of manufacture. We consider antibody “humanness,” which can be defined as the level of similarity between an observed antibody and commonly found antibodies sampled from humans, typically by sequencing the DNA of B-cells, which are responsible for producing antibodies. Similarity to antibodies from different sub-populations of B-cells may be more or less useful for different design purposes.

In this work, we explore that question by training specialized Naive B-cell models *AbBERT3 & AbBERT4*, in addition to more broadly-drawn collections of data *AbBERT1-2 & AbBERT5-9*. While other approaches to learning antibody properties exist, AbBERT provides an efficient solution by combining both heavy chains and light chains into a single training and prediction pipeline. This allows for a singular model to be robust enough to understand the nuances of different chain types. We also introspect on performance of various CDR regions through loop region annotation, leading to insights about mutational changes

### Related Works

ProtTrans (1) introduces auto-regressive and auto-encoder models trained on data from UniRef (2) and BFD (3). These models are able to capture biophysical features of protein sequences. We utilize the auto-encoder ProtBERT from this work for finetuning onto antibody sequences. Antibody humanization is also a closely related task which has been explored with BioPhi (4) where authors utilize Transformer models to implement a humanization platform. Most recently, AntiBERTa (5) is an approach to encoding B cell receptor (BCR) sequence representation utilizing attention-based language modeling.

## Data

The Observed Antibody Space (6)^1^ contains over 1 billion unpaired antibody sequences. This provides a rich data source for unsupervised learning of antibodies. Data was filtered by species, disease, and vaccine with options set to Human, None, and None, respectively. This was done in order to build out the required set for modeling human antibodies. By controlling for no disease and vaccine we can further control for purity in the human antibody with minimal outside influence. Collected sequences were then split into 200K, 2M, and 20M subsets. These subsets are evenly divided between heavy and light chains. An additional split of 2M and 20M was also performed with the added filter of BType assigned to Naive-B-Cells only.

### CDR Annotated Data

Due to their function as sensitive and specific identifiers of exogenous molecules, antibodies have several *hypervariable regions* of their sequence that largely determine binding behavior. These are typically referred to as the *complementarity determining regions* (CDRs); three lie in the heavy chain (‘H1’, ‘H2’, ‘H3’) and three in the light chain (‘L1’, ‘L2’, ‘L3’). For the purposes of antibody design, it is typically of greatest interest and difficulty to control the sensitivity and specificity of the antibody’s recognition of target molecules via modification of the CDRs. We thus explore using annotated sequence data, where the CDRs are provided to the language model. Models utilize the same data source splits defined above with the addition of “annotation tokens” inserted before and after each CDR. Annotation tokens were positioned according to Paratome, an automated antibody binding region identification tool (7). These annotation tokens are defined as additional tokens for the AbBERT tokenizer. The annotation tokens follow the notation of “START” and “L” or “H” for the chain type. Tokens are added to sequences by prepending the start token and appending the end token to a CDR region as for example, [S, S, [START_L1], Q, S…, T, Y [END_L1] …]. Antibody sequence annotation may be done in any of several, subtly different ways. Common approaches are North (8), IMGT (9), and ANARCI, (10). Our selection of Paratome is due to ease of use as well as the ability to automatically annotate H3, which is of particular interest for its binding properties and complexity. We defer further investigation of annotation methods and their relative merits.

## Methods

In this section we introduce our methods for learning and evaluating antibody sequences. Table 1 serves as a legend to our different experimental models and the unique setups associated with them. Here we introduce our approach to *“Chain Type”* token embeddings which take advantage of data augmentation through annotation tokens powered by CDR annotation tooling. Additionally we outline the procedure which is the basis for all downstream predictions known as “multi-mask” scoring.

**Table 1.** Available experimental model setups.

| MODEL | DATA SIZE | SOURCE | ANNO | FINETUNE |
| --- | --- | --- | --- | --- |
| AbBERT1 | 200K | HUMAN | ✓ | ✓ |
| AbBERT2 | 200K | HUMAN | × | ✓ |
| AbBERT7 | 200K | HUMAN | × | × |
| AbBERT5 | 2M | HUMAN | × | ✓ |
| AbBERT8 | 2M | HUMAN | × | × |
| AbBERT3 | 2M | NAIVE | × | ✓ |
| AbBERT6 | 20M | HUMAN | × | ✓ |
| AbBERT9 | 20M | HUMAN | × | × |
| AbBERT4 | 20M | NAIVE | × | ✓ |

### AbBERT Training Details

AbBERT experimental models were trained with 20k steps. Below we show the experimental setups to our models. The setups differ in their finetuning, data sources, annotations, and B-cell type (naive or human).

### “Chain Type” Token Embeddings

Chain type embeddings were included as part of the training procedure by passing a boolean matrix, 0’s for heavy chain input ids and 1’s for light chain input ids. This allows batching heavy and light chains during training time and is shown to be effective in

### Scoring

We implement a *multi-unmask* approach to scoring amino acid masking accuracy.

*Loop* regions (L1, L2, L3, or H1, H2, H3) are far more variable than the so-called “scaffold” and “constant” regions and so it is of special interest to see how our model performs on unmasking amino acids in only loop regions. Our primary motivation is to understand which modifications of the CDRs are more or less usual as an input to a sequence design or selection process. To further control results for loop regions only, masks for loop tokens are recorded during the annotation process as follows. We denote *l*_*ij*_ *∈* { 0, 1 } to be loop segment indicator variables, where 1 means the corresponding *x*_*ij*_ position belongs to a loop segment, and 0 is otherwise: *x*_*ij*_ belongs to a non-loop region. Further, we can expand *l*_*ij*_ from binary to categorical to denote which specific loop a token belongs to and then measure performance for these different loop regions. This expansion is the basis for our investigation on loop region H3.

The multi-unmasking validation scheme then goes as follows:

- A Bernoulli random variable *M*_*ij*_ is placed on each token *x*_*ij*_ in a sequence *S*_*i*_ with probability: *P* (*M*_*ij*_ = 1) = *p*, where we denote outcome 1 as the corresponding token being masked out and 0 as unmasked. Note that, following this scheme, amino acid tokens are being masked for prediction in an independent way. If some dependence or correlation is needed between the tokens, one would have to modify this masking distribution.
- If *loop_only_masking* is *True*, then we zero out the probability *p* for those *M*_*ij*_ where *l*_*ij*_ is 0. This means we do not mask out scaffold tokens and thus only amino acids in loop regions are masked out (independently) with probability *p*. Otherwise, when *loop_only_masking* is *F alse*, all tokens in both loop and scaffold regions have the same masking probabilities of *p*.
- The multi-unmasking scheme is defined below where tokens where *m*_*ij*_ == 1 (the realizations of *M*_*ij*_) are included in the computation of 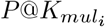. Formally, learning both chains in the same run. for each sequence *S*_*i*_:

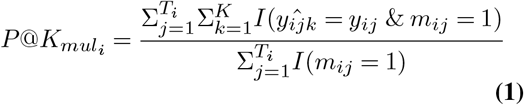

The results of this algorithm are visually illustrated in the experiments section below where *loop_only_masking is True*.

### *In Silico* Experiments

Using the methods described above we introduce experiments to evaluate the unmasking accuracy of the trained models as well as the ability to appropriately score therapeutic and immunogenic antibodies. The main objectives of the ummasking experiments is to show that the learning procedure is sufficient for learning common amino acid patterns. The scoring of therapeutics shows that data selection as well as training has sufficient information to perform on real world data. The results following are broken down into subsections of models as well as respective validation tasks. The first task “multi-unmask” follows the scoring laid out in section 4.3.1 and is performed on antibody sequences from Table 2. We focus on heavy chains as light chain unmasking is significantly easier even with models that have not been exposed to much data. Additionally, we pay special attention to hypervariable region 3 (H3) on the CDR for its interesting binding properties and complexity.

**Table 2.** Sequences picked as samples from design campaigns.

| ID | ANTIBODY | PDB |
| --- | --- | --- |
| 0 | M396 | 2G75 |
| 1 | s230 | 6NB8 |
| 2 | 80R | 2GHW |
| 3 | v2381 | — |
| 4 | v2130 | 7L7E |

#### Scoring Sample Sequences

We start out with investigating *AbBERT7* as our benchmark which shows the poorest accuracy on the masking task. Plots indicate unmasking accuracy averaged across *R*=20 runs with selected Top 1 accuracy. Error bars represent the standard deviation of the observations. Here the unmasking task is concerned with unmasking all loop regions (L1, L2, L3 for *light sample*) and (H1, H2, H3 for *heavy sample*)

**Fig. 1.**
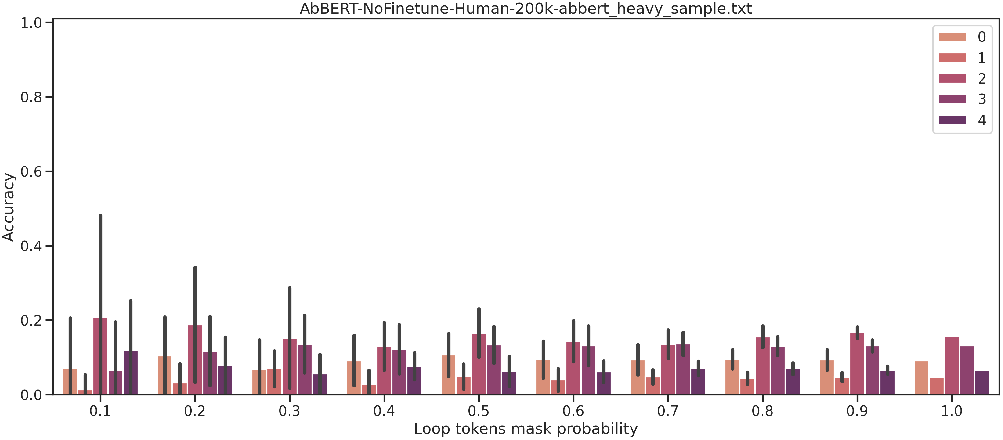
All loop region unmasking.

**Fig. 2.**
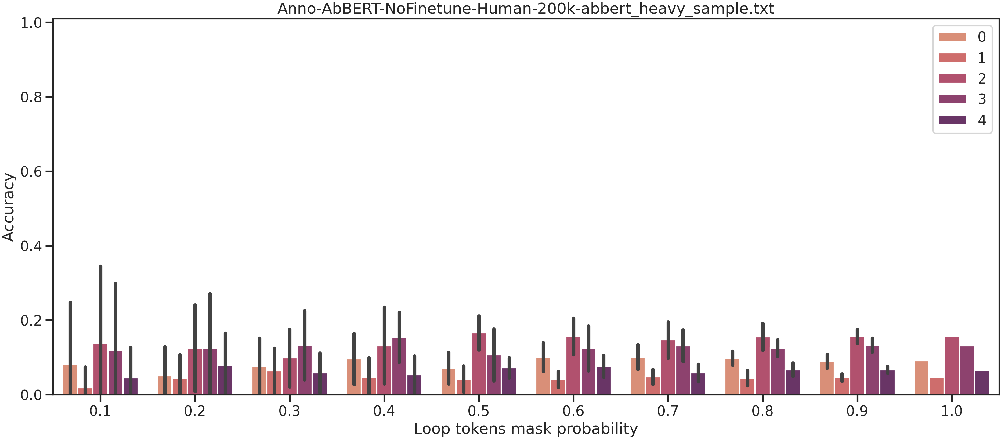
All loop region unmasking, annotated dataset.

Attempts were made to increase loop masking accuracy for the *AbBERT7* experiment by adding annotations to loop regions in attempts at providing context to loop regions during learning but yielded no significant improvements over the original dataset and in some cases performed worse. Expanding the dataset sizes to 2m and 20m sequences yielded marginal unmasking accuracy gains but was at best random for all loop mask tokens.

We then explore unmasking accuracy on H3 directly to establish baseline performance. We observe that models which are not finetuned are unable to achieve unmasking accuracy within any usable range for decision making. Furthermore we do not notice any trends with unmasking probabilities. That is, even with only 10 percent of an H3 region masked, there is no better accuracy than with 100 percent of the H3 region being masked. This holds true with annotated datasets as well as original sequences.

**Fig. 3.**
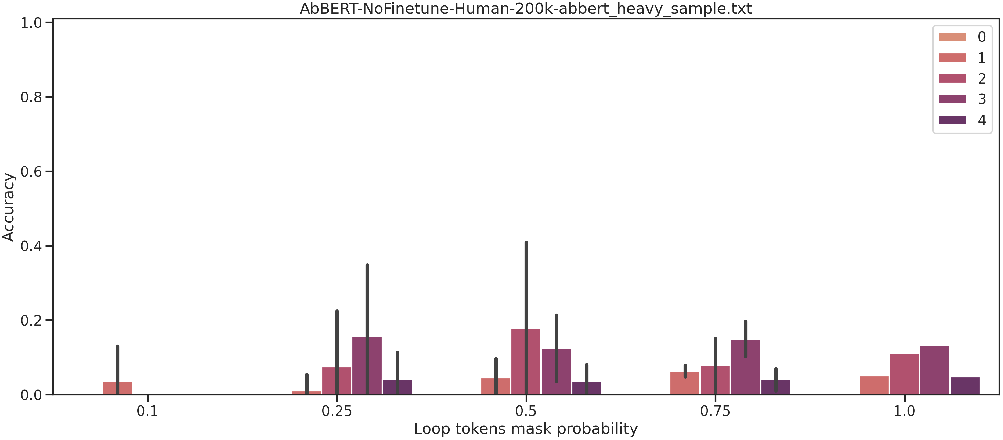
H3 unmasking probability no finetuning

**Fig. 4.**
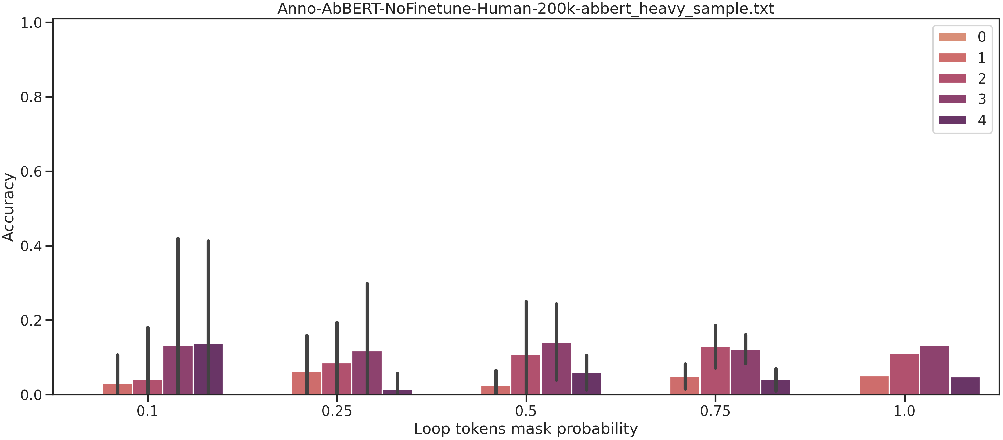
H3 unmasking probability no finetuning, annotated dataset.

Finetuned models see an increase in H3 unmasking accuracy when using annotated datasets. We actually see better performance on a smaller dataset of 200k unpaired sequences than when training AbBERT on 20M sequences. This may imply that as we see more diverse H3 regions but there is also an opportunity to explore annotation as applied to bigger datasets. Annotations prove to be beneficial in helping the model learn context for recovering H3 regions.

**Fig. 5.**
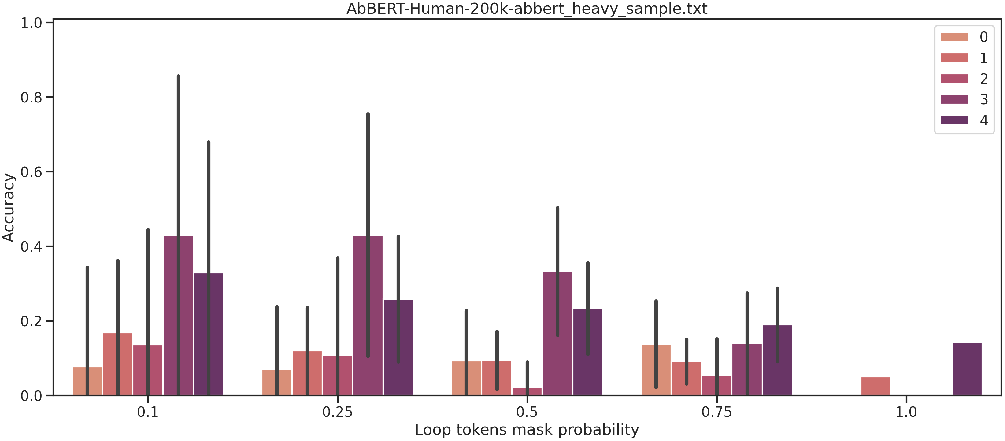
H3 unmasking probability with finetuning.

**Fig. 6.**
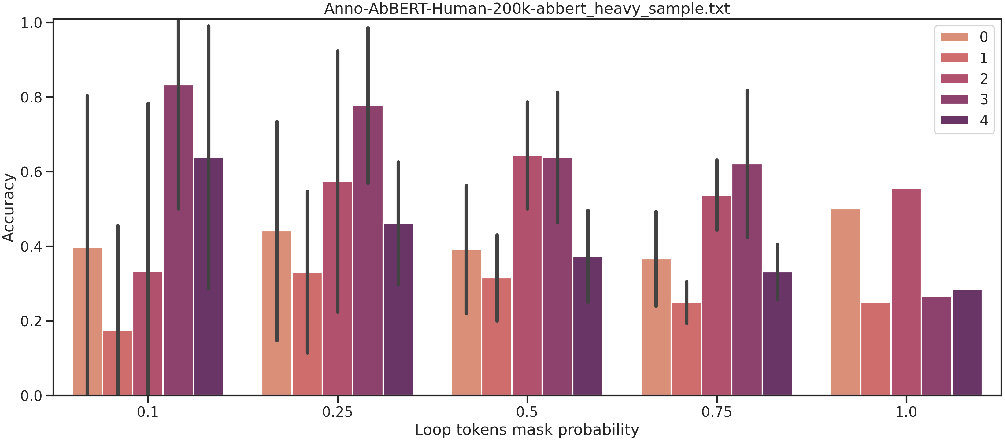
H3 unmasking probability with finetuning with annotated training data.

Next we attempt to further enhance our ability to recall H3 regions by utilizing Naive-B-Cell data from OAS as well as using larger sizes of data for human antibody sequences not specifically naive b cells. Our previous experiments prove annotations to be the leading drivers of successful H3 recall, strictly Naive-B-Cell sequence data learning slightly outperforms general human sequence data learning when unmasking region H3 even without annotations. This offers two viable approaches to understanding a notoriously complex region with interesting binding properties.

**Fig. 7.**
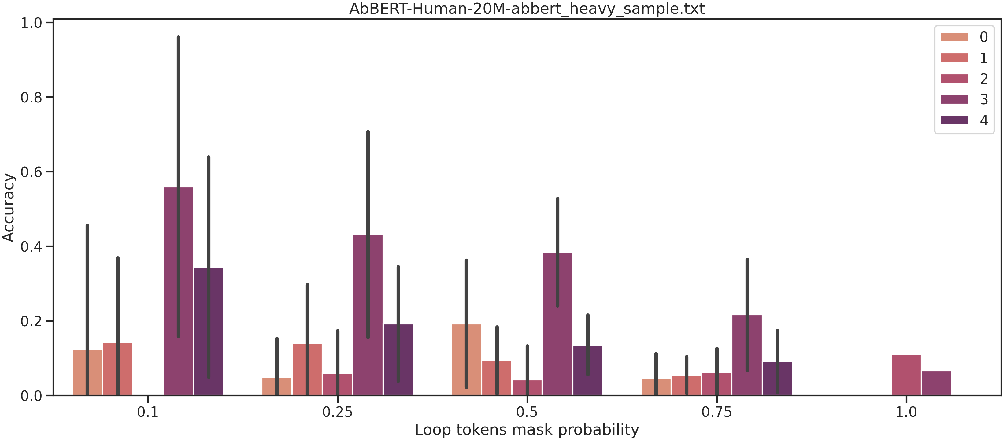
H3 unmasking probability with human training data.

#### Therapeutic Scoring

Therapeutic scoring evaluates how models perceive antibodies which have passed various clinical trials. For this task we selected over 600 therapeutic antibodies (11). While unmasking accuracy offers insights into masked language modeling learning performance it does not directly observe if what has been learned can be applied to real world design.

**Fig. 8.**
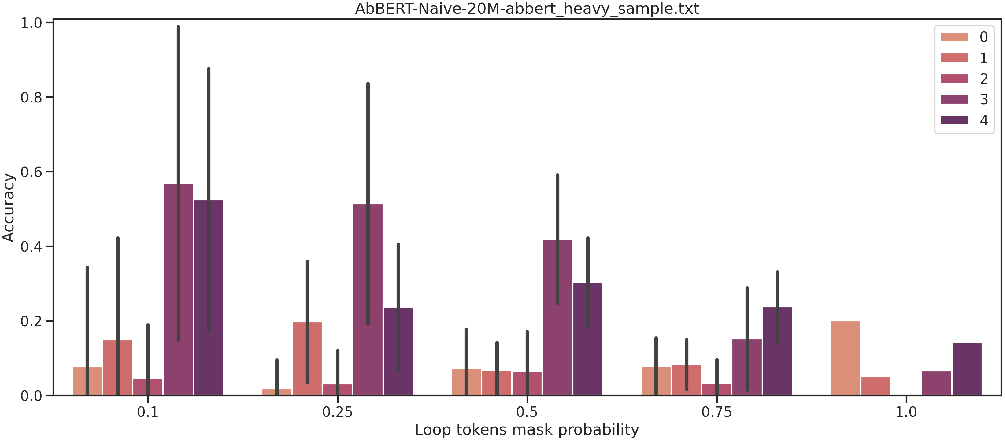
H3 unmasking probability with finetuning with naive B cell training data.

**Fig. 9.**
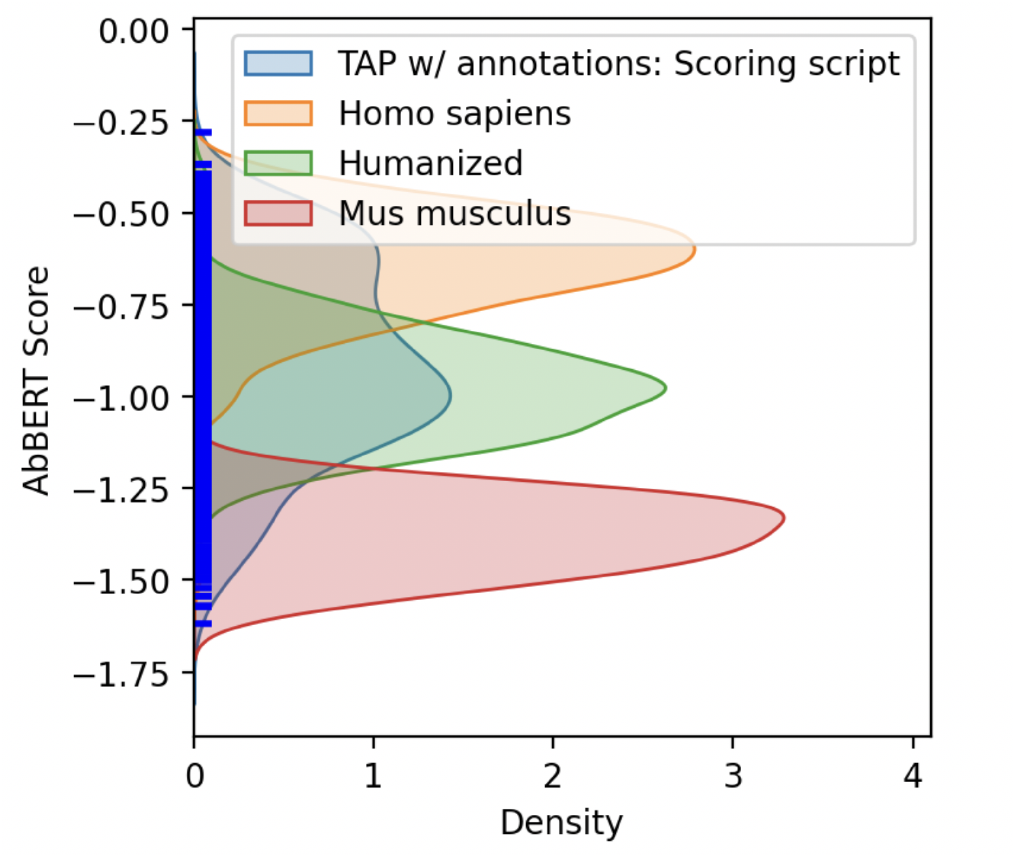
Therapeutics by species with Anno-AbBERT.

First we present two figures which illustrate the ability of models, both trained on datasets with and without annotation tokens to appropriately score distributions of therapeutics based on species. We see that Homo sapiens achieve the highest scoring. This implies the ability of AbBERT to learn proper characteristics of “humanness” from OAS data and through multi-unmask. Species such as Mus musculus (Rodent) are scored poorest and is to be expected as antibodies derived from these species often times diverge from typical human identified antibodies. We then observe where the distribution of therapuetics is shown as well as the distribution of immunogenic response antibodies (12). While there are some outliers the bigger distribution of immunogenics is scored considerably poorer than the therapeutic antibodies.

### Downstream Tasks Enabled by AbBERT

#### Deep Symbolic Optimization

In this section, we explore the *in silico* antibody optimization application, which employs simulation and optimization methods to identify promising antibody sequences that bind strongly to a target antigen.

**Fig. 10.**
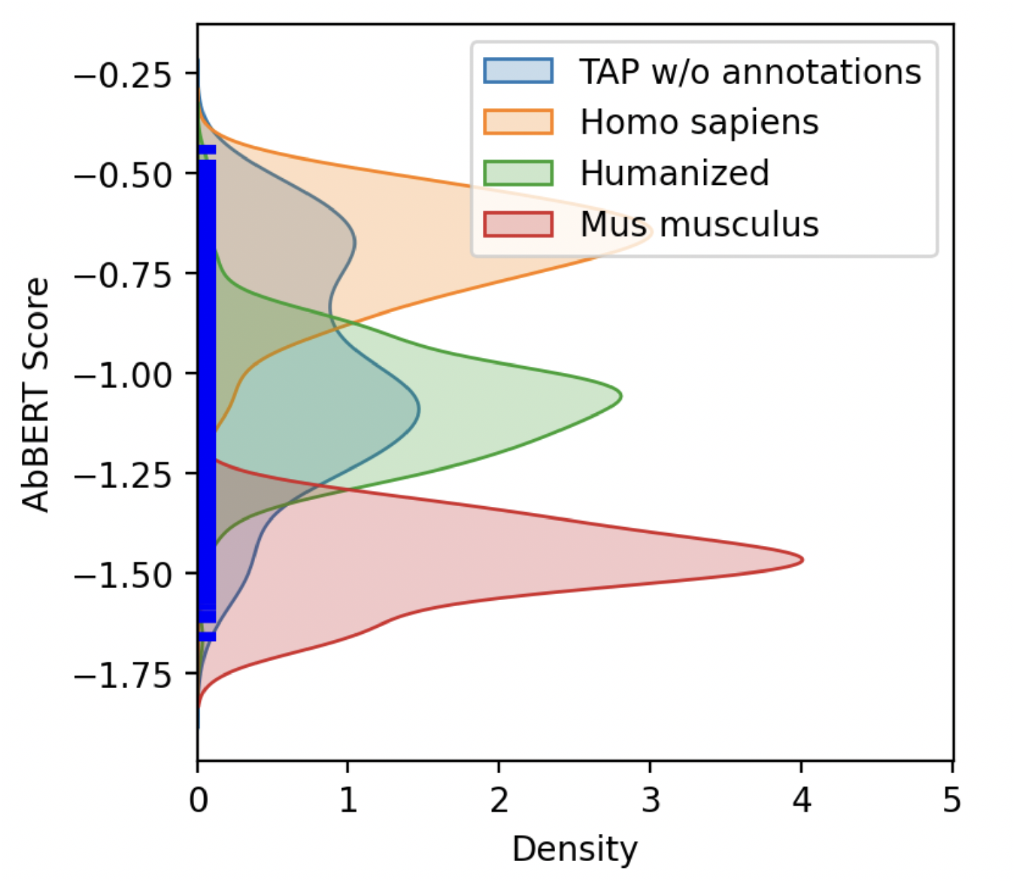
Therapeutics by species with AbBERT.

**Fig. 11.**
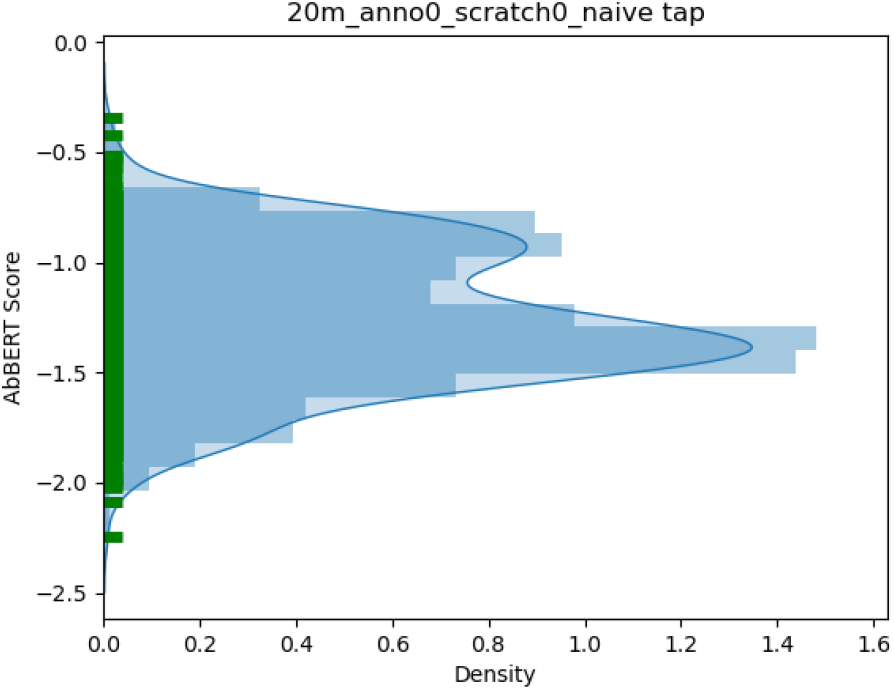
Therapeutics by species.

**Fig. 12.**
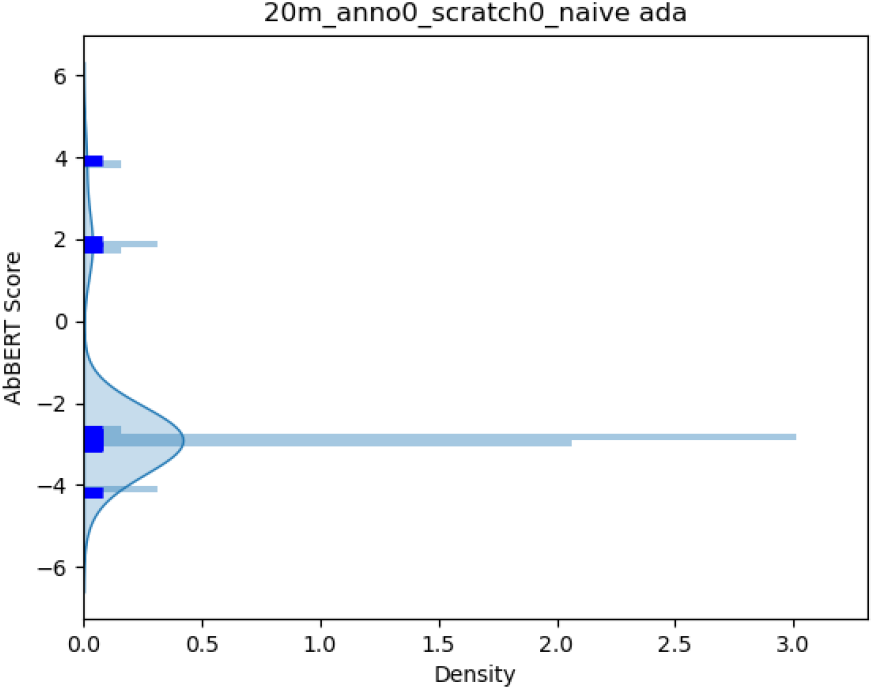
Immunogenics by species.

For this problem, we assume that a *base antibody* is available (in our case, an anti-SARS-CoV-1 antibody). This antibody is known to bind and neutralize a particular pathogen (SARS-CoV-1) but *not* the target pathogen (in our case, SARS-CoV-2). Our optimization task is to modify the amino acids comprising the CDRs of the base antibody to bind SARS-CoV-2. This can be performed by executing an optimizer that is able to replace amino acids from the base antibody in a way to optimize binding to the target pathogen. While classical search algorithms (such as genetic program methods) could in principle be employed in this problem, the difficulty of evaluating all possible combinations of amino acids (unconstrained search space in the order of 2040, while each single binding quality evaluation might take hours) makes it infeasible to apply any naive search strategy.

To solve this problem, we use the DSO (13) optimizer^2^, which we will refer to as *Baseline*. Although DSO solves the problem by sequentially sampling mutations in the base antibody and progressively learning which amino acids have to be replaced to optimize binding, it still takes a considerable time to find a new potentially antibody.

Part of the reason for that is because DSO does not have embedded knowledge of the relation between amino acids and has to learn how they interact initially by trial and error.

In order to accelerate this learning process, we provide AbBERT embeddings as additional observations to DSO, leveraging the knowledge about antibodies already embedded in AbBERT. This combination DSO+AbBERT enables solving the antibody optimization in an efficient way (a task that is not solvable by directly using a language model because there is no readily-available dataset of binding quality to our target pathogen).

For comparison purposes, we also evaluate DSO+ProtBERT

**Fig. 13.**
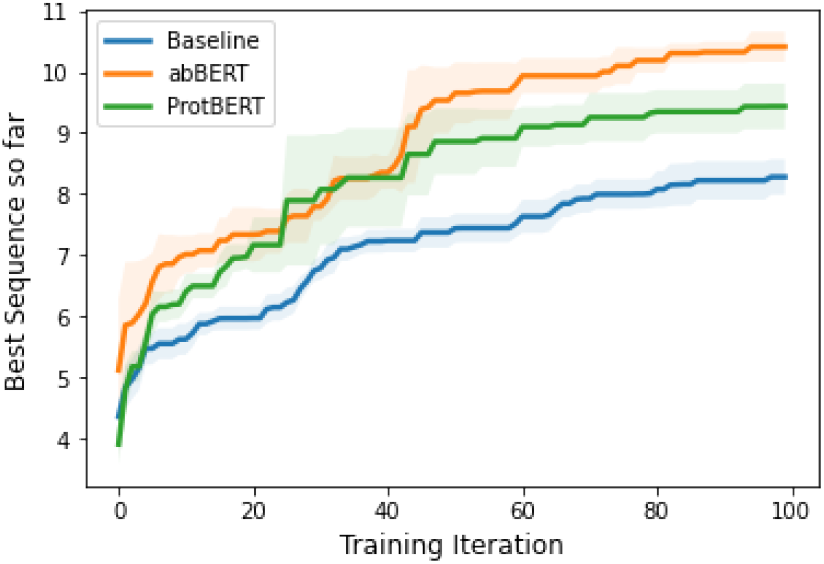
Quality of Best sequence found so far for the 80R antibody.

#### *In Vitro* Experiments

AbBERT has been used to support rapid antibody development with its scoring application. The following figures are drawn from an optimization campaign where the goal was to design effective antibodies against variants of SARS-CoV-2, starting with the COV2-2130 antibody (15).

**Fig. 14.**
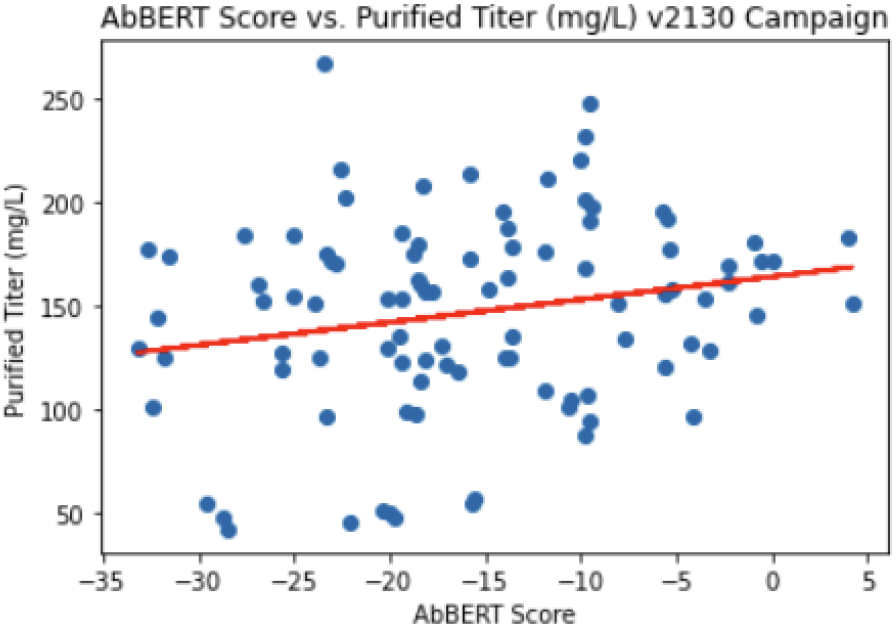
AbBERT vs. Purified Titer

We observe a positive correlation between the purified titer (the concentration of antibody protein produced) and AbBERT scoring for 96 exemplar antibodies. This result shows that those antibody sequences scored by AbBERT as more human-like tended to produce better, an important attribute in antibody design and selection.

Additionally, observe correlations with the AbBERT score and the predicted stability of the antibody via Free Energy Perturbation (16). While thermodynamic stability is not expected to be directly correlated with humanness, it may speak to different complementary aspects of antibody manufacturability tooling.

## Conclusions

AbBERT is a new approach towards modeling similarity of generated antibody sequences to human antibodies with an attention based language model. We have explored subsets of OAS data, primarily human sequence data with further specialized Naive-B-Cell data in attempts to learn about humanness properties. In addition we have applied CDR annotations in order to enhance the original data sets in hopes of learning difficult hypervariable region H3. We introduce chain type embeddings allowing our language model to batch heavy and light chains during learning as well as during scoring for antibody design campaigns simplifying the pipeline significantly. We illustrated that not only are we able to learn loop region unmasking effectively, we are able to increase our recall performance on region H3 through data annotation. These novel contributions lead us to positively scoring therapeutics and negatively scoring antibodies that are immunogenic. Finally we discuss a real world campaign to rapidly develop antibodies and show that AbBERT is successful in predicting poor manufacturability outcomes.

## ACKNOWLEDGEMENTS

The DOD’s Joint Program Executive Office for Chemical, Biological, Radiological and Nuclear Defense (JPEO-CBRND), in collaboration with the Defense Health Agency (DHA) COVID funding initiative for Rapid co-design of manufacturable and efficacious antibody therapeutics for COVID-19 via a machine-learning-driven computational design platform, molecular dynamics simulations, and experimental validation, Lawrence Livermore National Laboratory (LLNL), Proposal L22260, Agreement ID#44208 was used for this effort.

This work was performed under the auspices of the U.S. Department of Energy by Lawrence Livermore National Laboratory under Contract DE-AC52-07NA27344

LLNL-JRNL-838265

## Footnotes

1 OAS Dataset: http://opig.stats.ox.ac.uk/webapps/oas/oas

2 More information on the problem modeling and how the optimizer is set up in (14)

